# Novel surface plasmon resonance biosensor that uses full-length Det7 phage tail protein for rapid and selective detection of *Salmonella enterica* serovar Typhimurium

**DOI:** 10.1101/699488

**Authors:** Seok Hywan Hyeon, Woon Ki Lim, Hae Ja Shin

## Abstract

We report a novel surface plasmon resonance (SPR) biosensor that uses the full-length Det7 phage tail protein (Det7T) to rapidly and selectively detect *Salmonella enterica* serovar Typhimurium. Det7T, which was obtained using recombinant protein expression and purification in *Escherichia coli*, demonstrated a size of ∼75 kDa upon SDS-PAGE and was homotrimeric in its native structure. Micro-agglutination and TEM data revealed that the protein specifically bound to the host, *S.* Typhimurium, but not to non-host *E. coli* K-12 cells. The observed protein agglutination occurred over a concentration range of 1.5∼25 μg.ml^−1^. The Det7T proteins were immobilized on gold-coated surfaces using amine-coupling to generate a novel Det7T-functionalized SPR biosensor, wherein the specific binding of these proteins with bacteria was detected by SPR. We observed rapid detection of (∼ 20 min) and typical binding kinetics with *S.* Typhimurium in the range of 5 × 10^4^-5 × 10^7^ CFU.ml^−1^, but not with *E. coli* at any tested concentration, indicating that the sensor exhibited recognition specificity. Similar binding was observed with 10% apple juice spiked with *S.* Typhimurium, suggesting that this strategy could be expanded for the rapid and selective monitoring of target microorganisms in the environment.

## Introduction

Infections involving pathogenic bacteria are a major cause of morbidity and mortality in human beings and animals around world, resulting in a huge economic burden related to healthcare costs and manpower/wealth losses. Rapid monitoring of pathogenic microorganisms is critical for our ability to halt the fast spread of bacteria. The reliable conventional methods for detecting microorganisms (e.g., culture-based biochemical and serological assays) require the growth and technical manipulation of large amounts of cells, and thus tend to be time-consuming (24-52 h), labor-intensive, and cost-ineffective [1–3]. Biosensors are regarded as an attractive alternative, and have been used to detect various environmental pollutants [4–6] and microorganisms [7, 8]. A biosensor has two key components: a biological sensing element that enables specific recognition of a pathogen; and a transducer that converts this recognition to a measurable signal. Different sensing elements have been exploited in the development of biosensor platforms, including antibodies [9], DNA [10], RNA [11], aptamers [12], peptides [13], and carbohydrates [14]. Similarly, various transducing techniques have been reported, including optical [15], electrochemical [16, 17], mass perturbance-based [9], mechanical resonator-based [18], and SPR-based [19, 20] methods.

Bacteriophages have recently gained interest as sensing elements for pathogen-detecting platforms, because they are abundant in nature, stable under harsh conditions (e.g., extremes of temperature, pH, ionic strength, etc.), and specific to their target host bacteria [8]. Phages recognize their specific hosts through the ability of their tail proteins to bind receptors on the bacterial surface. The recognition and binding of a receptor by the tail protein is highly specific, which makes phages useful for bacterial typing and excellent candidates as sensing elements in biosensors [21]. To date, *E. coli* [22–25], *Staphylococcus aureus* [19, 20], and *Bacillus anthracis* spores [26] have been monitored using biosensors that include intact phages as recognition elements. However, intact phage-based detection has some limitations, such as loss of the biosensor signal due to lysis of the captured bacteria and a decrease in binding capacity due to the drying effect of long-term exposure [2]. Inconsistency in the signal arising from a biosensor platform may reflect the ability of some intact phages to enzymatically degrade surface receptors on their host bacteria. Moreover, intact phages are relatively large for the use of SPR, which is a distance-dependent sensing platform to detect refractive index change [8]. As an alternative strategy, the use of phage tail proteins has been proposed. Indeed, some phage tail proteins have been tested for bacterial detection [2, 21]. For example, cysteine-tagged P22 phage receptor binding proteins were used to detect *S.* Typhimurium [2] and a glutathione-S-transferase fusion protein from phage NCTC 12673 was used to detect *C. jejuni* [27, 28]. We recently showed that a fragment of tail protein from phage lambda (6HN-J) bound specifically to the host, *E. coli* K-12, but not to other bacteria [29]. We observed nonspecific transient attachment to non-host bacteria, and proposed that it might be part of the mechanism through which viral tail proteins recognize their host receptors. This was previously suggested by Silva et al. [30], who described that the first step of bacteriophage adsorption involves random collisions between the phage and various bacteria (e.g., by Brownian motion, dispersion, diffusion, and/or flow). During these interactions, the phage searches for its specific receptor via reversible (transient) binding [30].

In order to address the issues raised with intact phages or truncated tail proteins as a sensing element in biosensor, we examined whether the full-length tail protein of phage Det7 could be coupled with a biosensing platform to enable the specific recognition and capture of *S*. Typhimurium. Det7 is a *Salmonella* phage (*Myovirus*) whose 75-kDa tail protein exhibits 50% overall sequence identity to the tail endorhamnosidase of *Podovirus* P22 [31]. Both tail proteins bind octasaccharide fragments from *Salmonella* lipopolysaccharide, and that of Det7 was found to strongly infect all P22-susceptible strains of *Salmonella* and numerous additional *Salmonella* serovars, yielding a combined susceptibility of approximately 60% across all *Salmonella* strains [32]. Det7 is highly resistant to thermal unfolding and denaturation by SDS [31]. In the present study, we cloned and purified the full-length tail protein of Det7 (Det7T), then tested its specific binding to *S.* Typhimurium versus *E. coli* K-12 using micro-agglutination assays and TEM. Following this verification, we immobilized Det7T on gold substrates to form a novel SPR biosensor, then analyzed the host bacterial capture ability of the newly developed biosensor. Rapid detection and typical specific binding kinetics were observed against *S.* Typhimurium. The results indicate that the biosensor developed here shows the rapid, selective, and real-time monitoring of target microorganisms.

## Materials and methods

### Chemicals

Culture reagents were purchased from Difco (Detroit, MI). The plasmid isolation and gel extraction kits were obtained from Qiagen (Hilden, Germany). The pET Express & Purify kits (In-Fusion Ready) were from Clontech (Mountain View, CA). DEAE and Sephadex G-100 were purchased from Bio-Rad (Hercules, CA) and Sigma-Aldrich (Saint Louis, MO). The horseradish peroxidase (HRP)-conjugated monoclonal anti-His antibody (anti-mouse Cat. #631210) was obtained from Clontech (Mountain View, CA). The ultrasensitive HRP substrate used for Western blotting was from TaKaRa (Shiga, Japan). The CM5 Sensor Chip, N-hydroxyl succinimide (NHS), N-ethyl-N-dimethyl aminopropyl carbodiimide (EDC), ethanolamine, and PBS running buffer were purchased from BIAcore (GE Healthcare, Uppsala, Sweden). All other reagents used in the present work were of analytical grade and were applied without further purification.

### Bacteria and culture conditions

Plasmids were maintained in *E. coli* DH5α (TaKaRa, *hsdR, recA, thi-1, relA1, gyrA*96). *E. coli* K-12 and *S.* Typhimurium (x3339, Lab. of Bacterial Pathogenesis, Dept. of Microbiology, Pusan National University) were grown in tryptic soy broth at 37°C. The fusion protein was expressed in *E. coli* BL21 (DE3) (TaKaRa, *hsdS, gal, ëclts*857, *ind*1, *sam*7, *nin*5, *lac*UV5-T7gene1). Plasmid-containing *E. coli* DH5α and BL21 (DE3) cells were cultured at 37°C in TYS media (1% tryptone, 0.5% yeast extract, and 0.5% NaCl) containing 30 μ g.mL^−1^ ampicillin. Cells were stained with methylene blue and counted using a hemocytometer (Incyto Co., Cheonan, Korea) under a microscope (Zeiss Axiocam, Oberkochen, Germany).

### Cloning of Det7T

The full-length DNA encoding the tail protein from phage Det7 was PCR amplified with specific primers (forward 5’-AAG GCC TCT GTC GAC ATG ATT TCT CAA TTC AAT CAA-3’, reverse 5’-AGA ATT CGC AAG CTT TTA TTA CAC AGA TAA CTT CAT ACG-3’). The resulting PCR products were gel isolated, cloned into the N or C In-Fusion Ready vector (pET-6xHN-N, pET-6xHN-C) using an In-Fusion Ready cloning kit according to the manufacturer’s recommendations, and transformed into *E. coli* BL21 (DE3). The cells transformed with the desired plasmid (pDet7T) were identified by colony PCR. Cloning, ligation, and transformation were performed according to standard methods [33].

### Protein purification

*E. coli* BL21 (DE3) cells harboring pDet7T were incubated with isopropyl-β -D-thiogalactopyranoside (IPTG) to trigger overexpression of the fusion protein, which was purified using heat, DEAE, and Sephadex (G-100). Briefly, cells were cultured overnight, sub-cultured in 500 ml TYS-ampicillin, grown to OD _600 nm_ = 0.8, induced for 4 h with 1 mM IPTG, and harvested by centrifugation at 10,000 × g for 20 min. The cell pellets were frozen at −20°C and then suspended in a 1/10 volume (relative to the initial culture volume) of PBS buffer, and sonicated three times on ice (10 sec each with 30-sec pauses between bursts). The extract was centrifuged at 10,000 × g for 20 min at 4°C and the supernatant was transferred to a clean tube for SDS-PAGE analysis [29]. For purification, the supernatant was heated for 6 min at 80°C, centrifuged at 13,500 × g for 20 min at 4°C, and then applied to a DEAE column equilibrated with PBS. The fractions were collected by elution with a NaCl linear gradient (0-400 mM) in PBS. Those containing Det7T were pooled, further purified using a Sephadex G-100 column, and collected for additional analyses.

### Western blot analysis

Soluble and purified protein fractions were subjected to SDS-PAGE and transferred to nitrocellulose membranes using standard techniques [33]. Each membrane was incubated for 1 h at room temperature with 20 ml blocking buffer (5% nonfat dry milk in 0.2% Tween-20/PBS), and then incubated (with shaking) for 1 h at room temperature with horseradish peroxidase (HRP)-conjugated monoclonal anti-His antibody (diluted 1:4000 in blocking buffer)[29]. Each membrane was washed three times (10 min per wash) with washing buffer (0.2% Tween-20 in PBS), incubated (with shaking) with a chemiluminescent HRP substrate for 5 min, and exposed to an X-ray film [29].

### Micro-agglutination assay

The micro-agglutination assay [27] was performed with Det7T and bacteria in microtiter plates at 4°C. Overnight cells (OD_600 nm_ = 1) were centrifuged at 10,000 × g for 5 min and suspended with 1 ml PBS. *S.* Typhimurium or control *E. coli* cells (50 μl per well of the above suspensions) were mixed with two-fold serial dilutions of Det7T in PBS (0.024∼25 μ g.ml^−1^) and the plates were incubated overnight at 4°C. Wells in which the cells appeared diffused were taken as containing agglutinated cells, whereas wells that had round dots (sedimented cells) at their bottoms were taken as containing non-agglutinated cells.

### TEM

Overnight-cultured cells (approximately 5 × 10^7^ cells.ml^−1^) were centrifuged 10,000 × g for 5 min and suspended with 1 ml PBS. A Formvar-coated copper grid was sequentially placed on a 1:1 mixture of 0.05 M NHS and 0.2 M EDC for 20 min, purified Det7T (∼25 μg.ml^−1^) for 30 min, PBS for 5 min (3 times), and 1 M ethanolamine (pH 8.5) for 15 min. After the excess active sites were activated, immobilized, and deactivated, the grid was washed several times with PBS, exposed to the indicated microorganism for 30 min, washed three times with PBS, dried, and observed by TEM (Hitachi H-7600, Tokyo, Japan) [29].

### Surface Plasmon Resonance (SPR)

The binding kinetics of Det7T with the microorganisms were studied using a BIAcore T200 instrument (GE Healthcare, Little Chalfont, UK). Det7T was diluted (1:50) in running buffer (PBS) and immobilized on a Series S CM5 Chip using a standard amine-capture method. The two cells of the chip were treated separately during the immobilization procedure: flow cell 1 was spared the attachment of Det7T and used as a blank reference surface, while flow cell 2 was coupled with Det7T. The chip was primed with running buffer (PBS), activated by a flow of 0.4 M EDC/0.1 M NHS mixture (1:1) for 420 sec, immobilized with Det7T (1:50 diluted, 1.14 mg.ml^−1^) in immobilization buffer (10 mM sodium acetate, pH 4.0) for 180 sec, and deactivated by 1 M ethanolamine (pH 8.5) for 420 sec (S1 Fig). The chip was then assessed with different numbers of microorganisms (heat-killed in boiling water for 15 min) for 20 min. Serial dilutions of the microorganism were prepared in running buffer or 10% apple juice in running buffer (for environmental sample assessments) and flowed through the cells. Binding analysis was conducted at a flow rate of 5 μl.min^−1^ at 25°C throughout the experiment, with the exception of the regeneration step (30 μl.min^−1^). The sensor surface was regenerated for 20 sec using 10 mM NaOH. In each run, the association phase and subsequent dissociation phase were each monitored for 10 min. The binding sensorgrams were estimated for each microorganism using the obtained reference-subtracted sensorgrams (representing that of flow cell 2 minus that of flow cell 1) and a steady-state affinity model, as applied using the Biacore T200 evaluation software (version 3.0) [29].

## Results and discussion

### Overexpression, purification, and Western blot analysis of Det7T

The Det7 phage appears to offer advantages over other viral sensing elements for *Salmonella* detection, as it has a wide-ranging host specificity against various *Salmonella* sp. [32], and its tail protein exhibits good heat stability [31]. Here, we set out to use the full-length wild-type Det7 tail protein (Det7T) as a sensing element for SPR biosensor construction. A construct encoding full-length Det7T was PCR amplified to yield a product of ∼2130 base pairs. We fused the purified PCR product with a (His)_6_-tag by inserting it into a linearized vector in which the coding sequence was under the control of the IPTG-inducible T7*lac* promoter. In our system, Det7T was found to be overproduced in the soluble fraction, as assessed by SDS-PAGE (Fig 1A).

**Fig 1.**
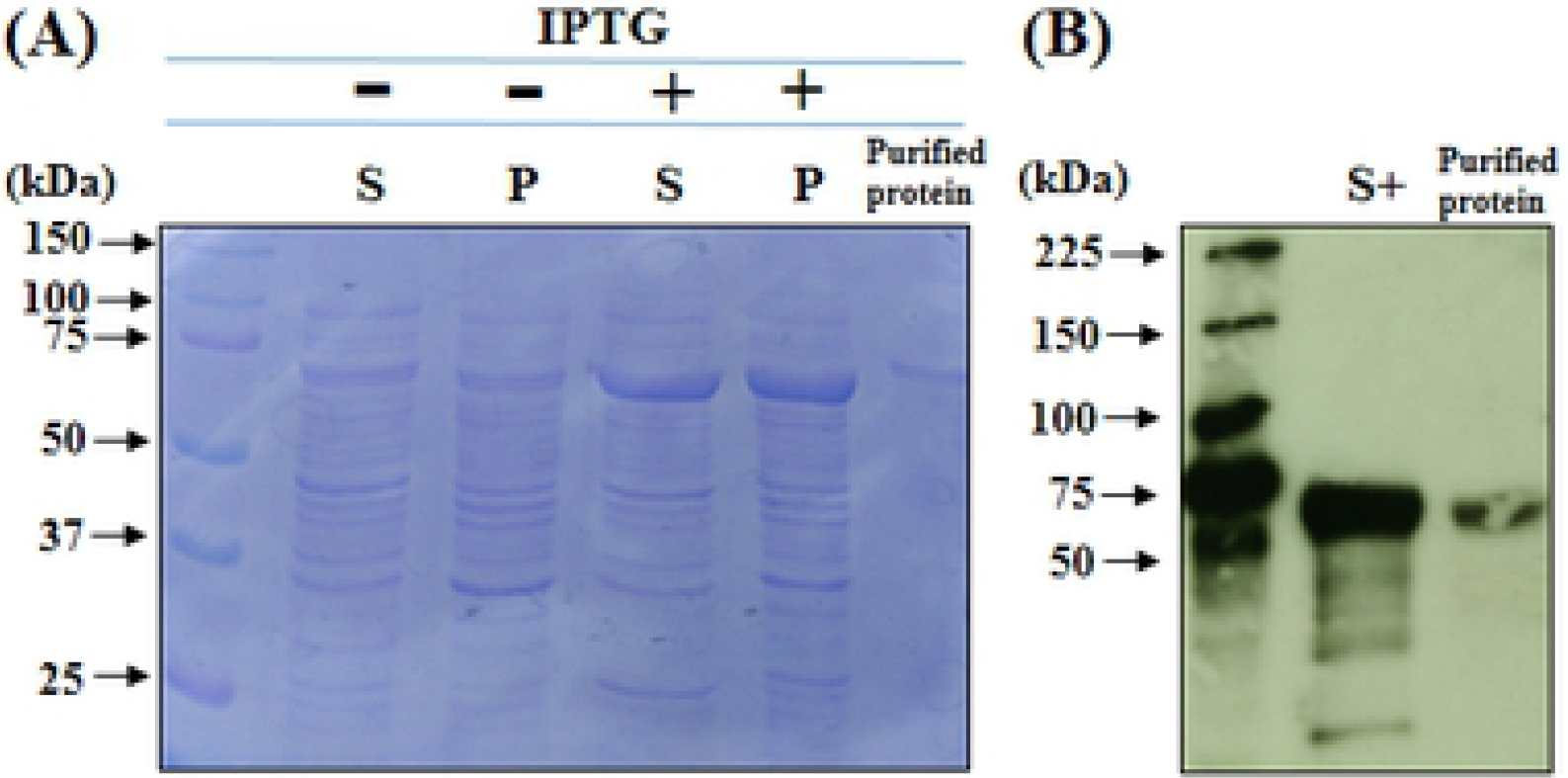
SDS-PAGE (A) and Western blotting (B) of the Det7 tail protein. (A) *E. coli* cells harboring pDet7T were induced with (lanes 2 and 3) or without (lanes 4 and 5) 1 mM IPTG. The cells were lysed and centrifuged to obtain pellet (P) and soluble (S) fractions. The purified Det7T (lane 6) had an apparent molecular weight of ∼75 kDa, as assessed using 12% SDS-PAGE. Lane 1, molecular size markers. (B) The resolved proteins were electro-transferred to a nitrocellulose membrane and incubated with an HRP-conjugated anti-His monoclonal antibody (1:4,000). Lane 2, soluble fraction obtained following induction with IPTG; lane 3, purified Det7T.

Although we initially applied this soluble fraction to a HisTALLON affinity column, we found that Det7T did not bind to this Ni column. This may reflect that the His-tagged N- or C-termini of Det7T are buried in its 3D structure, even under the slightly denaturing conditions (6 M urea) used in our work. It has been reported that the surface area of the monomer was buried in the trimer of the amino-terminally shortened Det7 tail protein, Det7tspΔ 1-151 [31]. Therefore, we heat treated the soluble fraction at 80°C for 6 min to denature other proteins, and then purified our target protein using DEAE and a Sephadex column. The purified Det7T had an apparent size of ∼75 kDa on SDS-PAGE (Fig 1A), which was consistent with the predicted size.

The identity of purified Det7T was confirmed by Western blotting with an anti-His monoclonal antibody, which clearly bound to the overexpressed protein band in the IPTG-induced soluble fraction (Fig 1B, lane 2) and the purified Det7T fraction (Fig 1B, lane 3), after both were boiled with sample loading buffer. Together, our results indicate that we obtained overexpressed and purified proteins corresponding to the expected (His)_6_-tagged fusion proteins. We thus set out to test the potential for the relatively small Det7T to be used for constructing an SPR-based biosensor.

### Micro-agglutination assay

The ability of the purified Det7T to bind *S.* Typhimurium or *E. coli* was tested by a micro-agglutination assay (Fig 2).

**Fig 2.**
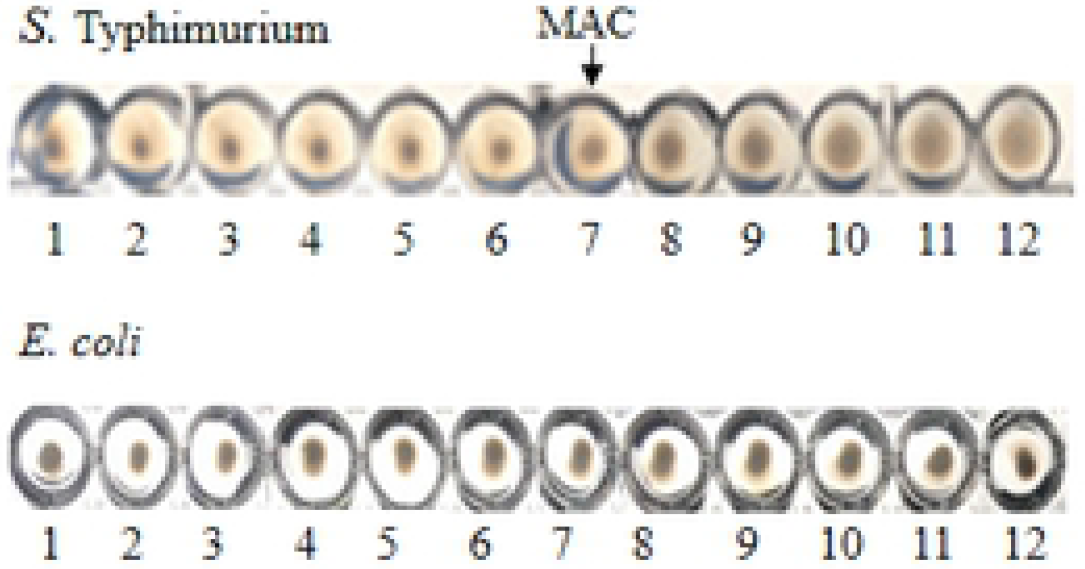
Micro-agglutination assay. Overnight cultures of *S.* Typhimurium or *E. coli* K-12 cells were washed with PBS and incubated with 2-fold serially diluted Det7T (0.024∼25 µg.ml^−1^) overnight at 4°C. Det7T agglutinates *S.* Typhimurium effectively across a concentration range of 1.5-25 μg.ml^−1^, as shown by a diffuse cell pattern at the bottom of wells (7^th^-12^th^ wells of the upper panel), while no agglutination was observed with *E. coli* K-12 (lower panel), even at the highest tested concentration. MAC: minimum agglutination concentration.

Given that Det7T is homotrimeric and binds to the surface of host cells [31], we assumed that it might cross-link and agglutinate with host *S.* Typhimurium cells. In a micro-agglutination assay, non-agglutinated bacteria form a compact sediment at the bottom of a well, while agglutinated bacteria form diffuse aggregates. Micro-agglutination was performed by incubating a constant number of bacterial cells with 2-fold dilutions (0.024∼25 μg.ml^−1^) of Det7T at 4°C. The diffuse pattern first appeared at the bottom of microplate wells in which 1.5 μg.ml^−1^ of Det7T was incubated with *S.* Typhimurium cells (Fig. 2, 7^th^ well on left of the upper panel); the degree of agglutination increased up to ∼25 μg.ml^−1^ (Fig 2) and plateaued thereafter. These results indicate that Det7T formed a tight bond with *S.* Typhimurium cells. The minimum concentration of Det7T that yielded detectable cell agglutination (1.5 μg.ml^−1^) was 2-fold lower than the minimal agglutinating concentration of P22 tail protein (3 μg.ml^−1^) [27]. There was no detectable specific binding of Det7T with *E. coli* (Fig 2), indicating that Det7T specifically agglutinated with *S.* Typhimurium.

### TEM

The binding of Det7T to the surface of *S.* Typhimurium cells or *E. coli* (control) was visualized with TEM. A Formvar-coated grid was incubated sequentially on the NHS/EDC mixture, Det7T (∼25 μg.ml^−1^), and ethanolamine for activation, immobilization, and deactivation of excess active groups respectively. The grid was then washed several times with PBS, exposed to the microorganism, and observed by TEM. Consistent with the results of our micro-agglutination assays, we observed the binding of Det7T to *S.* Typhimurium cells treated with Det7T (Fig 3A), but not to Det7T-treated *E. coli* (Fig 3B) or to non-treated *S.* Typhimurium cells (Fig 3C).

**Fig 3.**
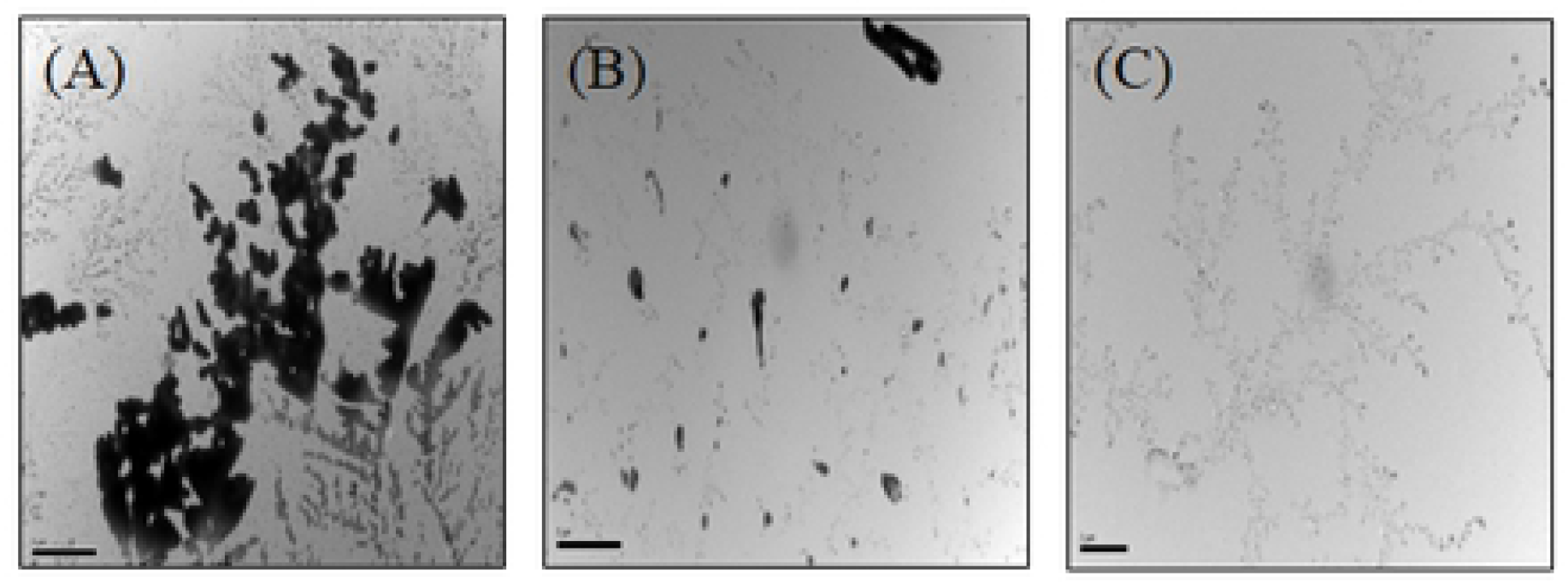
Electron microscopic study of Det7T. A Formvar-coated copper grid was sequentially placed on NHS/EDC (1:1) for 20 min, Det7T (∼ 25 μg.ml^−1^) for 30 min, PBS for 5 min (3 times), 1M ethanolamine (pH 8.5) for 15 min, PBS for 5 min (three times), and *S.* Typhimurium (A) or *E. coli* K-12 (B) cells for 30 min. The grid was then washed several times with PBS, dried, and observed by TEM (Hitachi H-7600). “Blank” indicates *S.* Typhimurium without Det7T treatment (C).

This result is in agreement with our recent finding that binding of anti-rabbit immunoglobulin-gold was observed in host *E. coli* K-12 cells pretreated with purified 6HN-J protein, but not non-pretreated K-12 cells or non-host *Pseudomonas aeruginosa* [29].

### SPR

The binding kinetics of Det7T with host *S.* Typhimurium or non-host *E. coli* were assessed using an SPR instrument (BIAcore T200) with a Series S CM5 Chip. Flow cell 1 lacked Det7T and was used as blank reference; this enabled us to use the Biacore T200 evaluation software (version 3.0) to subtract the nonspecific bindings of microbes from the obtained sensorgrams. We obtained higher responses when the sensor surface was coupled with ∼1.14 μ g.ml^−1^ Det7T (1:50 dilution, ∼560 RU). We thus used 1:50-diluted Det7T our binding assays. The sensorgrams of live and heat-killed microbes were very similar, but the former showed slight oscillations; thus, we used heat-killed microbes for our binding assays. The change in SPR angle obtained with various bacterial concentrations was taken as indicating the interaction of bound Det7T with bacteria. Typical binding kinetics (association and dissociation) were obtained with *S.* Typhimurium cells in the range of 5 × 10^4^-5 × 10^7^ CFU.ml^−1^ (Fig 4A).

**Fig 4.**
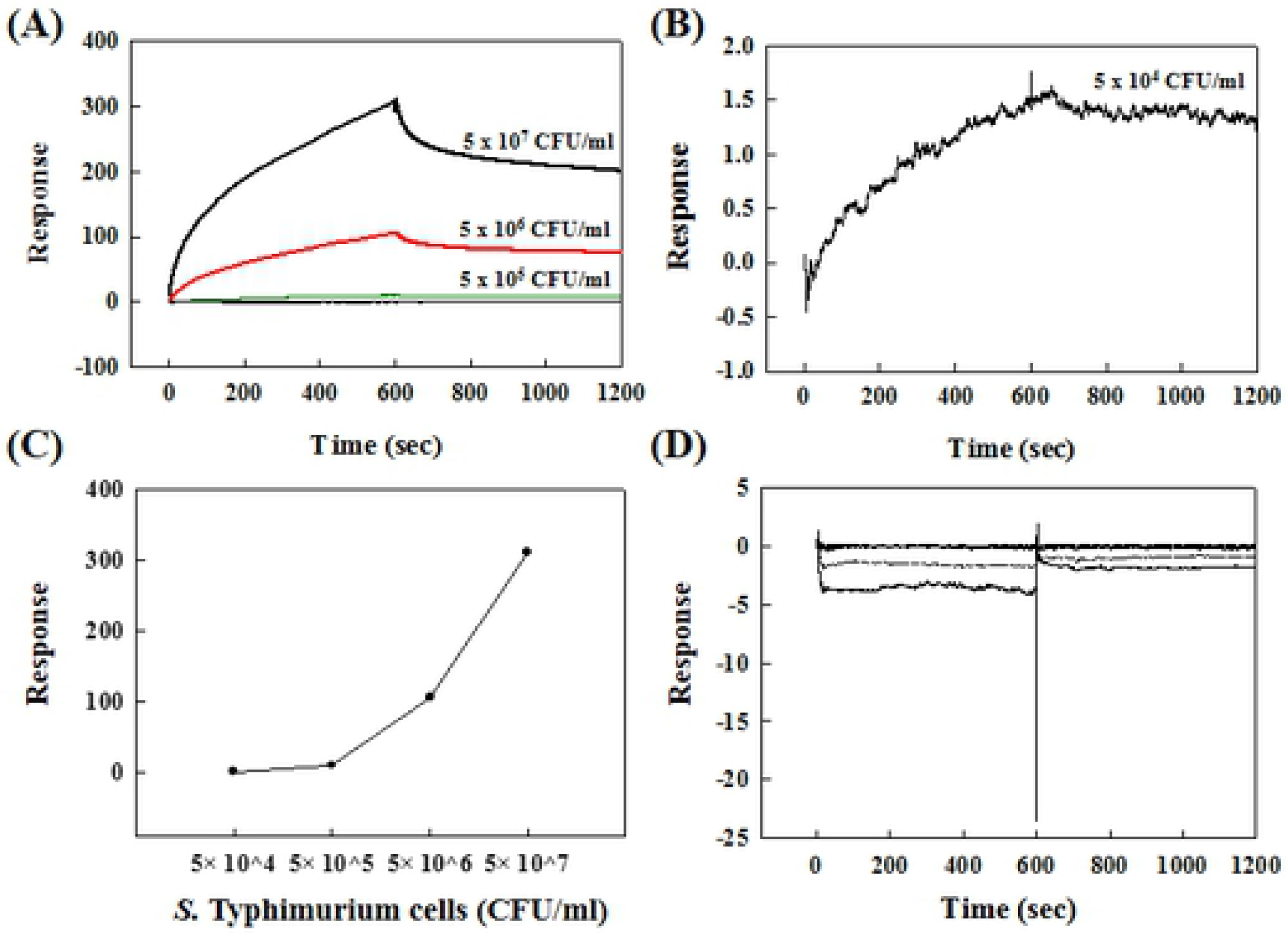
SPR responses of Det7T. (A) Det7T (1:50, 1.14 mg.ml^−1^) was attached to the CM5 chip for 1 min, and different amounts (5 × 10^4^-5 × 10^7^ CFU.ml^−1^) of *S.* Typhimurium cells were flowed over the sensor surface for 20 min at a flow rate of 5 μl.min^−1^. (B) The response obtained with *S.* Typhimurium cells at 5 × 10^4^ CFU.ml^−1^was re-plotted with a different y-scale. (C) A dose-dependent response was observed. (D) *E. coli* K-12 (2 × 10^4^-2 × 10^7^ CFU.ml^−1^) cells were flowed for 20 min under the same conditions.

We obtained a significant signal of bacterial capture with as little as ∼5 × 10^4^ CFU.ml^−1^ *S*. Typhimurium (Fig 4B). We previously obtained a comparable detection limit of 2 × 10^4^ CFU.ml^−1^ for *E. coli* K-12 with immobilization of N-terminally (His)_6_-tagged 6HN-J on a Ni-coupled chip [29]. Our results resemble those of a report in which the detection limit of *S. aureus* was found to be 10^4^ CFU.ml^−1^ with the lytic phage SPR-based SPREETA^TM^ sensor [19]. The SPR response representing *S.* Typhimurium binding was found to be dose-dependent (Fig 4C). In contrast, we did not observe any binding response to non-host *E. coli* (Fig 4D). Sensorgrams for *E. coli* demonstrated negative responses below those obtained with the reference, indicating that Det7T did not recognize lipopolysaccharide on the surface of *E. coli* as a specific binding receptor. These SPR data are quite different from those obtained in our previous study involving a truncated 6HN-J protein, which transiently attached and detached from the surface of non-host bacteria (*P. aeruginosa*) while binding firmly to the surface of host *E. coli* [29]. Based on these previous findings, we had speculated that the phage tail protein might use transient reversible attachment to a cell surface as a means to search for a specific receptor to initiate phage translocation, as mentioned by Silva’s group [30]. Here, we found that full-length Det7T showed rapid recognition and binding to the cell surface of host cells (*S.* Typhimurium), but no response (even transient attachment) with non-host cells (*E. coli*). Therefore, we propose a new hypothesis for our previous findings: We speculate that 6HN-J, which is a fragment of tail protein, might lack the domain(s) involved in forming a tight and specific bond with its receptor on the bacterial surface. Another possibility is that Det7T can quickly recognize and bind to the outermost lipopolysaccharide of its host, whereas 6HN-J might need time to search for and recognize its surface receptor, LamB.

Lastly, we tested whether the Det7T-containing biosensor could detect *S.* Typhimurium in an environmental sample. A 10% solution of apple juice was spiked with *S.* Typhimurium and applied to the SPR biosensor system. As shown in Figure 5, the Det7T-based biosensor recognized and bound to the spiked *S.* Typhimurium cells, yielding sufficient signals across concentrations of 5 × 10^5^ to 5 × 10^7^ CFU.ml^−1^.

**Figure 5.**
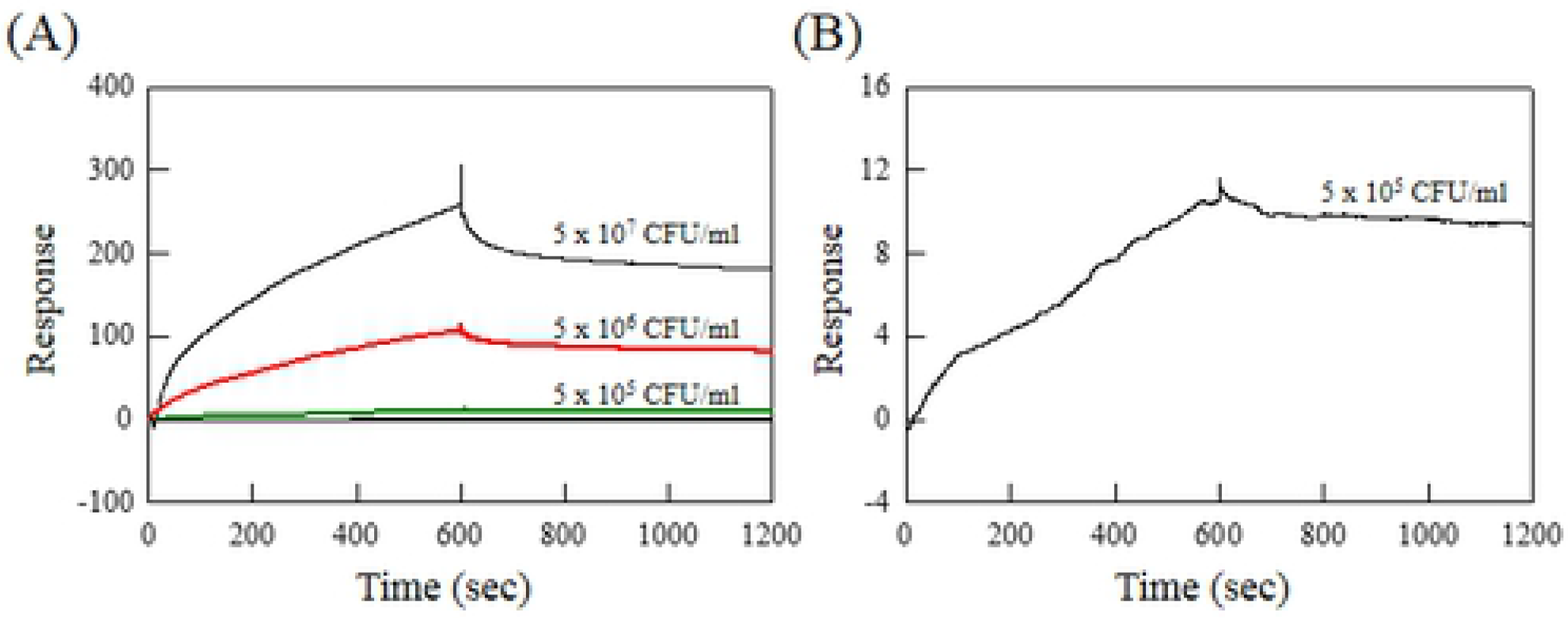
Detection of microorganisms in apple juice using the SPR biosensor. (A) Different amounts (5 × 10^5^-5 × 10^7^ CFU.ml^−1^) of *S.* Typhimurium cells were spiked into 10% apple juice and flowed over the sensor surface at a flow rate of 5 μl.min^−1^. (B) The response obtained with *S.* Typhimurium cells of 5 × 10^5^ CFU.ml^−1^ was re-plotted with a different y-scale.

These data suggest that the SPR biosensor system described herein could be exploited for rapidly and selectively monitoring pathogenic microbes in food and the environment.

## Conclusions

We herein exploited the full-length tail protein from phage Det7 as a sensing element for the SPR-based detection of *S.* Typhimurium. The purified full-length Det7T exhibited a size of 75 kDa upon SDS-PAGE and showed specific binding to host *S.* Typhimurium but not non-host *E. coli* K-12, as assessed by micro-agglutination and TEM. We then bound the generated Det7T to a CM5 chip through amine coupling to generate a novel Det7T-functionalized SPR biosensor. We observed rapid detection of (∼ 20 min) and typical binding kinetics with host *S.* Typhimurium in the range of 5 × 10^4^-5 × 10^7^ CFU.ml^−1^, but not with non-host *E. coli*, indicating that our biosensor exhibited selective recognition. The binding of Det7T was also similarly observed in 10% apple juice spiked with *S.* Typhimurium. The results suggest that our newly developed biosensor could be expanded for the rapid, selective, and real-time monitoring of target microorganisms.

## Supporting information

**S1 Fig. CM5 chip treatment using a standard amine-capture method.**

## Acknowledgements

The authors thank Dr. Nina K. Bröker of Universität Potsdam for providing the plasmid encoding the tail protein of phage Det7. We are grateful to Un Joo Gang for help with SPR and data analysis.

## Author Contributions

**Conceptualization:** Hae Ja Shin

**Funding acquisition:** Hae Ja Shin

**Investigation:** Hae Ja Shin, Seok Hywan Hyeon

**Methodology:** Hae Ja Shin, Seok Hywan Hyeon

**Supervision:** Hae Ja Shin, Woon Ki Lim

**Validation:** Hae Ja Shin, Seok Hywan Hyeon, Woon Ki Lim

**Writing – original draft:** Hae Ja Shin

**Writing – review & editing:** Woon Ki Lim

